# Rare modification in the ergosterol biosynthesis pathway leads to amphotericin B resistance in *Candida auris* clinical isolates

**DOI:** 10.1101/2021.10.22.465535

**Authors:** Milena Kordalewska, Kevin D. Guerrero, Timothy D. Mikulski, Tony N. Elias, Rocio Garcia-Rubio, Indira Berrio, Dianne Gardam, Christopher H. Heath, Anuradha Chowdhary, Nelesh P. Govender, David S. Perlin

## Abstract

We determined amphotericin B (AmB) susceptibility and sequenced key genes of the ergosterol biosynthesis pathway implicated in AmB resistance (*ERG2, ERG3, ERG6, ERG11*) of 321 clinical isolates of *Candida auris*. In antifungal susceptibility testing, 19 (5.9%) isolates were categorized as AmB-resistant (MIC ≥2 mg/l). Only one AmB-resistant isolate presented a unique non-wild-type *ERG6* genotype that was confirmed to confer amphotericin B resistance (MIC >32 mg/l) when introduced into a susceptible strain (MIC = 0.5 mg/l).

---

Amphotericin B (AmB), a polyene antifungal drug, has a broad-spectrum activity against pathogenic fungi, including *Candida* spp. (1). Its mode of action is rather unusual, as it does not inhibit an enzyme. Instead, AmB binds to ergosterol (ERG) (2), an abundant sterol that regulates permeability and fluidity of the fungal cell membrane (3). In most *Candida* species, AmB resistance is rare in comparison to resistance to other antifungal drug classes (azoles and echinocandins) (4). Most often, AmB resistance stems from alterations in the sterol composition of the fungal cell membrane due to mutations in genes of the ERG biosynthesis pathway. Whole genome sequence (WGS) analysis of AmB-resistant clinical isolates revealed a transposon insertion in *ERG2* (C-8 sterol isomerase) of *C. albicans*, while a missense mutation in *ERG3* (C-5 sterol desaturase) and deletion of 170 nucleotides in *ERG11* (lanosterol 14α-demethylase) were detected in *C. tropicalis* (4). Moreover, a single AmB-resistant *C. albicans* isolate was reported to harbor a substitution in *ERG11* and a sequence repetition (10 duplicated amino acids) leading to loss of function of *ERG5* (C-22 sterol desaturase) (5). In *C. glabrata* clinical isolates, targeted gene sequencing revealed the presence of AmB resistance-conferring mutations in *ERG2* and *ERG6* (C-24 sterol methyltransferase) (6-8). Additionally, *in vitro* evolution experiments confirmed the role of *ERG6* mutations in AmB resistance in *C. albicans* (4).

Clinical isolates of *C. auris*, a recently emerged nosocomial pathogen, were reported to have a higher prevalence of AmB resistance (based on tentative breakpoints), e.g., 30% in the U.S. (9). In contrast to relatively well-studied mechanisms of azole and echinocandin resistance (10-16), information about *C. auris* response to AmB is scarce. So far, suggestions for an underlying mechanism of AmB resistance in *C. auris* have come from WGS analysis. The researchers did not find any loss-of-function mutations in genes previously implicated in AmB resistance in *C. albicans* but listed various genes with non-synonymous mutations which may potentially play a role (17, 18). However, no follow-up studies (e.g., targeted gene sequencing, genetic engineering) were reported to confirm these observations.

The current antifungal armamentarium is extremely limited, with only three classes of systemic drugs widely available to treat *Candida* spp. infections. Antifungal treatment of *C. auris* infections is further complicated by several factors, including the scale of azole resistance (e.g. 90% of *C. auris* isolates in the U.S. are fluconazole-resistant) (9), emergence of multidrug resistance involving two or more drug classes (19, 20), and scarce availability of first-line therapy drugs, echinocandins, in resource-limited countries (21). Thus, a better understanding of the scale and molecular mechanism of AmB resistance in *C. auris* is urgently needed.

Here, we analyzed distribution of AmB minimal inhibitory concentration (MIC) values for 321 clinical isolates of *C. auris* representing five clades (I – South Asian, n=48; II – East Asian, n=6; III – South African, n=30; IV – South American, n=236; V – Iranian, n=1). Antifungal susceptibility testing (AFST) with AmB was performed with Etest gradient diffusion strips (bioMérieux, Marcy-l’Étoile, France) according to the manufacturer’s instructions. As recommended by the Centers for Disease Control and Prevention (CDC), MICs of 1.5 mg/l were rounded up to 2 mg/l. A tentative AmB MIC breakpoint of ≥2 mg/l, determined by the CDC on the basis of pharmacokinetic/pharmacodynamic study results (murine model of *C. auris* infection), was used to categorize isolates as resistant to AmB (9). A summary of AFST results and MIC values for individual isolates are presented in Table 1 and Supplementary Table 1, respectively. Three hundred and two isolates (94.1%) exhibited AmB MIC values <2 mg/l and were categorized as AmB-susceptible. A total of 19 isolates (5.9%), including 3 isolates of clade I, 1 isolate of clade III, and 15 isolates of clade IV, exhibited AmB MIC values ≥2 mg/l and therefore were categorized as AmB-resistant (Table 1). This rate is similar to the one reported recently from South Africa (6%) (16), but considerably lower than majority of the studies conducted so far, where anywhere from 10% to 35% of isolates (30% of U.S. isolates according to the CDC) were reported as AmB-resistant (9, 22-24).

**Table 1.** Results of AFST and gene sequencing performed for 321 *C. auris* isolates belonging to five geographic clades: I – South Asian (n=48); II – East Asian (n=6); III – South African (n=30); IV – South American (n=236); V – Iranian (n=1). AmB – amphotericin B; MIC – minimal inhibitory concentration * The actual range is 1.5-2 mg/l, but all AmB MIC values of 1.5 mg/l obtained by Etest were rounded up to 2, as recommended by the CDC. ** K177R/N335S/E343D – present in all isolates of clade IV

| Clade | Number of isolates | AmB MIC range [mg/l] | Erg2 | Erg3 | Erg6 | Erg11 |
| --- | --- | --- | --- | --- | --- | --- |
| **I** | 45 | 0.19 - 1 | WT | WT | WT | WT; Y132F; K143R |
|  | 3 (6.3%) | 2* | WT | WT | WT | Y132F |
| **II** | 6 | 0.094 - 0.5 | D39E | WT | WT | WT |
|  | 0 | ≥2 | - | - | - | - |
| **III** | 29 | 0.19 - 0.38 | D39E | WT | WT | V125A/F126L |
|  | 1 (3.3%) | 6 | D39E | WT | deletion of AA 18-181 | V125A/F126L |
| **IV** | 221 | 0.25 - 1 | D39E | S58T | WT | WT**; E102K; Y132F; K143R; G459S; I466M; Y501H |
|  | 15 (6.4%) | 2* | D39E | S58T | WT | WT** |
| **V** | 1 | 0.19 | WT | I31V/S37N/L63M | A287T | K177R/N335S/E343D/T366I |
|  | 0 | ≥2 | - | - | - | - |

To decipher the molecular resistance mechanism in isolates exhibiting elevated MIC values (≥2 mg/l), we amplified and sequenced the following genes of ergosterol biosynthesis pathway (in the entire collection of clinical isolates): *ERG2* (C-8 sterol isomerase; B9J08_004943), *ERG3* (C-5 sterol desaturase; B9J08_003737), *ERG6* (sterol 24-C-methyltransferase; B9J08_005340), and *ERG11* (lanosterol 14-α-demethylase; B9J08_001448), which were previously implicated in AmB resistance in other *Candida* spp. Gene sequences of *C. auris* strain B8441 extracted from FungiDB (fungidb.org) served as a reference for primer design (Supplementary Material 2) and sequence analysis. Primers were synthesized by Integrated DNA Technologies (Coralville, IA, United States), and Sanger sequencing was performed by Genewiz (South Plainfield, NJ, United States). A summary of sequencing results and genotypes of individual isolates are presented in Table 1 and Supplementary Table 1, respectively. In 18 of 19 AmB-resistant isolates no mutations in *ERG2, ERG3, ERG6*, or *ERG11* were found that could explain elevated AmB values (Table 1). We noticed that amplification of *ERG6* from one South African isolate (SA18, clade III; AmB MIC = 6 mg/l) yielded a much shorter PCR product (Figure 1). Sequence analysis revealed that this isolate presented a unique non-wild-type *ERG6* genotype where 492 base pairs (from 52 to 543) were deleted (SA18’s *ERG6* is 636 bp long in comparison to the wild-type’s (WT’s) 1128 bp), which corresponds to the deletion of 164 amino acids (from 18 to 181) and a shorter Erg6 (SA18’s Erg6 is 211 amino acids long in comparison to the 375 AA of WT) (Figure 2). Representative sequences were deposited at GeneBank (NCBI) with the accession numbers OK564516 - OK564551 (*ERG2*), OK564552 - OK564587 *(ERG3*), OK564588 - OK564623 (*ERG6*), and OK564624 - OK564654 (*ERG11*). Accession numbers for individual isolates are presented in Supplementary Table 1.

**Figure 1.**
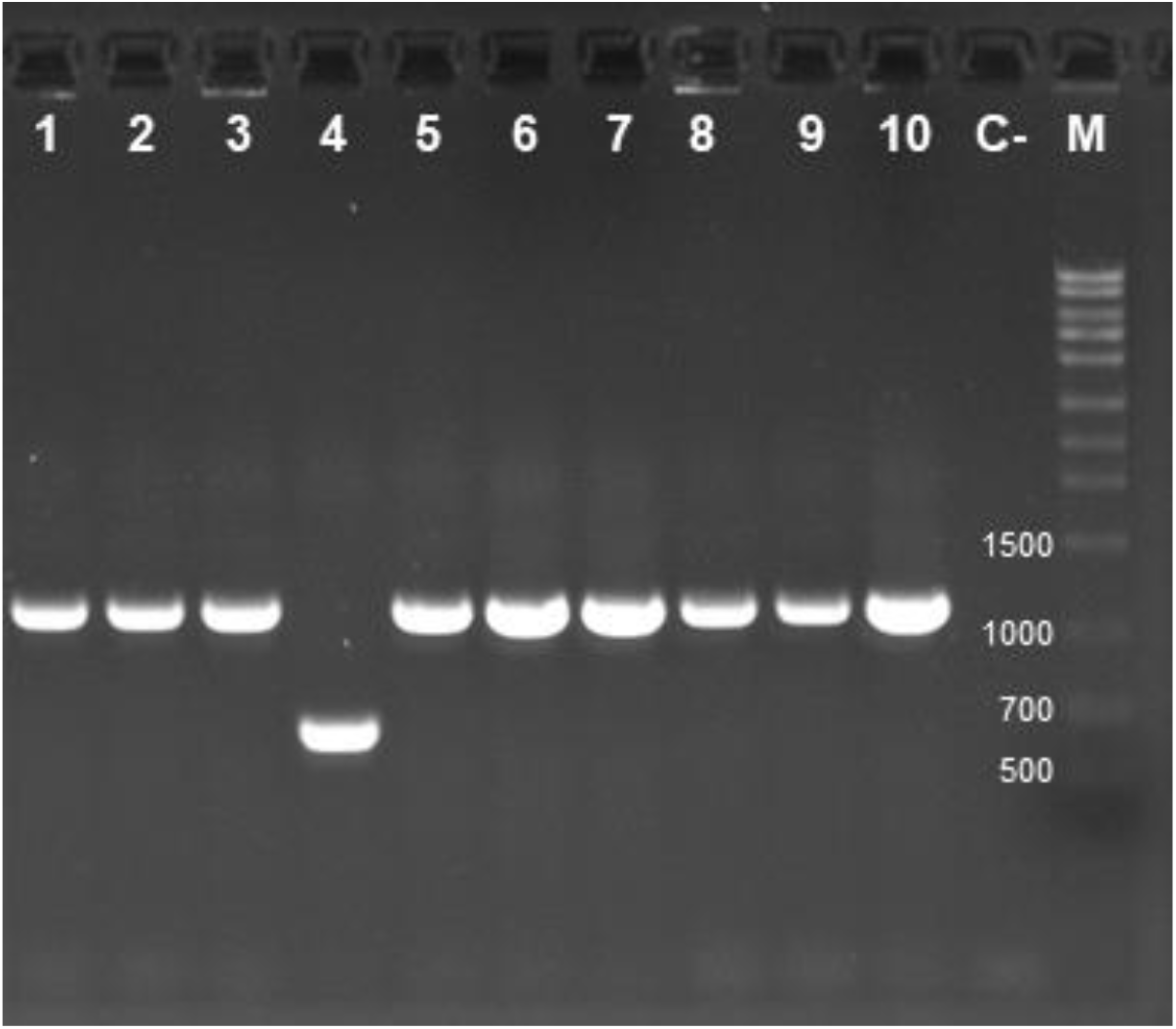
Results of PCR amplification of *ERG6* gene from *C. auris* clinical isolates SA15-SA24 (lanes 1-10). C-, negative control; M: molecular size marker

**Figure 2.**
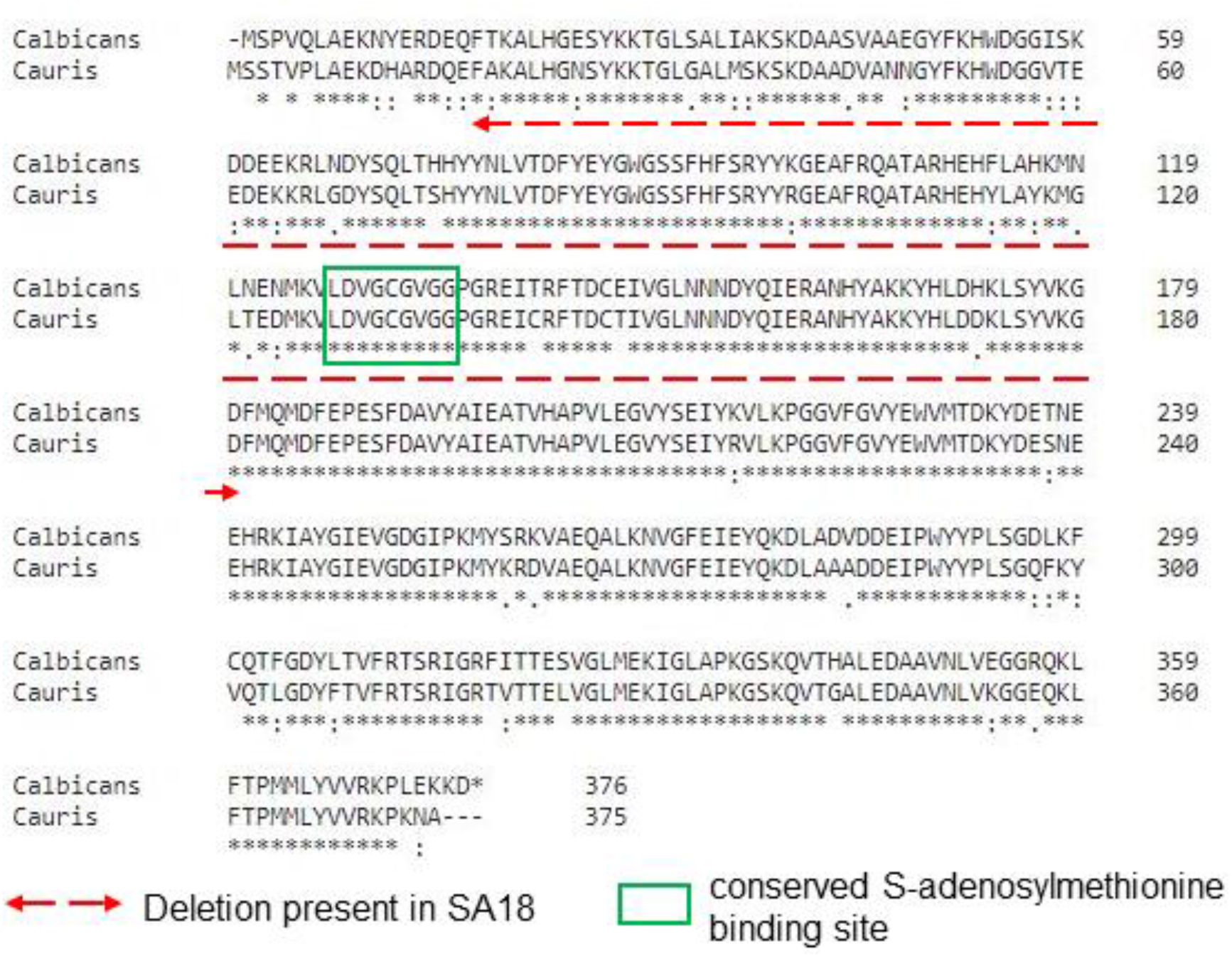
Alignment of Erg6 sequence from *Candida albicans* and *C. auris*.

To confirm that the SA18’s *ERG6* variant confers AmB resistance, a WT *ERG6* gene in AmB-susceptible (MIC = 0.5 mg/l) isolate VPCI 717/P/14 (clade I) was replaced with an *ERG6* of SA18 fused with nourseothricin (NAT) resistance gene by using CRISPR/Cas9 system as described before (25), except that the cells were made electrocompetent with the Frozen-EZ yeast transformation kit (Zymo Research, Irvine, CA, USA). The transformants were selected on YPD plates containing 300 mg/l NAT. Correctness of transformation was validated by PCR and sequencing. All PCR conditions, reagents and primers used are listed in Supplementary Material 2. After that, the AmB MIC of the correct transformant (MKKG066) was determined by Etest. MIC >32 mg/l (no zone of inhibition around the Etest strip) was read, indicating that amphotericin B resistance was induced when a wild-type *ERG6* was replaced with *ERG6* of SA18 in a susceptible strain.

*ERG6* encodes C-24 methyltransferase, which converts zymosterol to fecosterol in the ERG biosynthesis pathway. In *C. albicans*, a fragment between amino acids 127 and 135 of Erg6 is a highly conserved S-adenosylmethionine binding site (26). This fragment (amino acids 128-136 in Erg6 of *C. auris*) is not present in SA18’s Erg6 due to the deletion of amino acids 18-181 (Figure 2). It was previously shown that lack of Erg6 activity leads to the accumulation of zymosterol, which can support fungal cell growth, but the absence of ergosterol in the cell membrane confers AmB resistance (6).

In conclusion, we found that mutations in key genes of ergosterol biosynthesis, which can be linked directly to AmB resistance, are extremely rare. Only 1 of 313 clinical isolates screened (0.3%) had an *ERG6* variant which induced AmB resistance in a WT strain. It is possible that mechanisms other than *ERG6* mutations may also contribute to reduced AmB susceptibility in *C. auris*, although this remains to be determined.

## ACKNOWLEDGEMENTS

D.S.P. receives funding from the U.S. National Institutes of Health (NIH) and contracts with Merck, Regeneron and Pfizer. He serves on advisory boards for Amplyx, Astellas, Cidara, Matinas, N8 Medical and Scynexis. N.P.G. receives funding from the NIH, the U.S. Centers for Disease Control and Prevention (CDC), the U.K. Medical Research Council (MRC), and Gates Foundation. The remaining authors declare that the research was conducted in the absence of any commercial or financial relationships that could be construed as a potential conflict of interest.

## REFERENCES

1. Chandrasekar P. 2011. Management of invasive fungal infections: a role for polyenes. J Antimicrob Chemother 66:457–65.

2. Carolus H, Pierson S, Lagrou K, Van Dijck P. 2020. Amphotericin B and Other Polyenes-Discovery, Clinical Use, Mode of Action and Drug Resistance. J Fungi (Basel) 6.

3. Douglas LM, Konopka JB. 2014. Fungal membrane organization: the eisosome concept. Annu Rev Microbiol 68:377–93.

4. Vincent BM, Lancaster AK, Scherz-Shouval R, Whitesell L, Lindquist S. 2013. Fitness trade-offs restrict the evolution of resistance to amphotericin B. PLoS Biol 11:e1001692.

5. Martel CM, Parker JE, Bader O, Weig M, Gross U, Warrilow AG, Kelly DE, Kelly SL. 2010. A clinical isolate of Candida albicans with mutations in ERG11 (encoding sterol 14alpha-demethylase) and ERG5 (encoding C22 desaturase) is cross resistant to azoles and amphotericin B. Antimicrob Agents Chemother 54:3578–83.

6. Ahmad S, Joseph L, Parker JE, Asadzadeh M, Kelly SL, Meis JF, Khan Z. 2019. ERG6 and ERG2 Are Major Targets Conferring Reduced Susceptibility to Amphotericin B in Clinical Candida glabrata Isolates in Kuwait. Antimicrob Agents Chemother 63.

7. Vandeputte P, Tronchin G, Larcher G, Ernoult E, Berges T, Chabasse D, Bouchara JP. 2008. A nonsense mutation in the ERG6 gene leads to reduced susceptibility to polyenes in a clinical isolate of Candida glabrata. Antimicrob Agents Chemother 52:3701–9.

8. Vandeputte P, Tronchin G, Berges T, Hennequin C, Chabasse D, Bouchara JP. 2007. Reduced susceptibility to polyenes associated with a missense mutation in the ERG6 gene in a clinical isolate of Candida glabrata with pseudohyphal growth. Antimicrob Agents Chemother 51:982–90.

9. Centers for Disease Control and Prevention. 05/29/2021. Antifungal Susceptibility Testing and Interpretation. https://www.cdc.gov/fungal/candida-auris/c-auris-antifungal.html. Accessed 08/03/2021.

10. Chowdhary A, Prakash A, Sharma C, Kordalewska M, Kumar A, Sarma S, Tarai B, Singh A, Upadhyaya G, Upadhyay S, Yadav P, Singh PK, Khillan V, Sachdeva N, Perlin DS, Meis JF. 2018. A multicentre study of antifungal susceptibility patterns among 350 Candida auris isolates (2009-17) in India: role of the ERG11 and FKS1 genes in azole and echinocandin resistance. J Antimicrob Chemother 73:891–899.

11. Healey KR, Kordalewska M, Jimenez Ortigosa C, Singh A, Berrio I, Chowdhary A, Perlin DS. 2018. Limited ERG11 Mutations Identified in Isolates of Candida auris Directly Contribute to Reduced Azole Susceptibility. Antimicrob Agents Chemother 62.

12. Kordalewska M, Lee A, Park S, Berrio I, Chowdhary A, Zhao Y, Perlin DS. 2018. Understanding Echinocandin Resistance in the Emerging Pathogen Candida auris. Antimicrob Agents Chemother 62.

13. Rybak JM, Doorley LA, Nishimoto AT, Barker KS, Palmer GE, Rogers PD. 2019. Abrogation of Triazole Resistance upon Deletion of CDR1 in a Clinical Isolate of Candida auris. Antimicrob Agents Chemother 63.

14. Rybak JM, Munoz JF, Barker KS, Parker JE, Esquivel BD, Berkow EL, Lockhart SR, Gade L, Palmer GE, White TC, Kelly SL, Cuomo CA, Rogers PD. 2020. Mutations in TAC1B: a Novel Genetic Determinant of Clinical Fluconazole Resistance in Candida auris. mBio 11.

15. Li J, Coste AT, Liechti M, Bachmann D, Sanglard D, Lamoth F. 2021. Novel ERG11 and TAC1b mutations associated with azole resistance in Candida auris. Antimicrob Agents Chemother doi:10.1128/AAC.02663-20.

16. Maphanga TG, Naicker SD, Kwenda S, Munoz JF, van Schalkwyk E, Wadula J, Nana T, Ismail A, Coetzee J, Govind C, Mtshali PS, Mpembe RS, Govender NP, for G-S. 2021. In Vitro Antifungal Resistance of Candida auris Isolates from Bloodstream Infections, South Africa. Antimicrob Agents Chemother 65:e0051721.

17. Escandon P, Chow NA, Caceres DH, Gade L, Berkow EL, Armstrong P, Rivera S, Misas E, Duarte C, Moulton-Meissner H, Welsh RM, Parra C, Pescador LA, Villalobos N, Salcedo S, Berrio I, Varon C, Espinosa-Bode A, Lockhart SR, Jackson BR, Litvintseva AP, Beltran M, Chiller TM. 2019. Molecular Epidemiology of Candida auris in Colombia Reveals a Highly Related, Countrywide Colonization With Regional Patterns in Amphotericin B Resistance. Clin Infect Dis 68:15–21.

18. Munoz JF, Gade L, Chow NA, Loparev VN, Juieng P, Berkow EL, Farrer RA, Litvintseva AP, Cuomo CA. 2018. Genomic insights into multidrug-resistance, mating and virulence in Candida auris and related emerging species. Nat Commun 9:5346.

19. Lyman M, Forsberg K, Reuben J, Dang T, Free R, Seagle EE, Sexton DJ, Soda E, Jones H, Hawkins D, Anderson A, Bassett J, Lockhart SR, Merengwa E, Iyengar P, Jackson BR, Chiller T. 2021. Notes from the Field: Transmission of Pan-Resistant and Echinocandin-Resistant Candida auris in Health Care Facilities - Texas and the District of Columbia, January-April 2021. MMWR Morb Mortal Wkly Rep 70:1022–1023.

20. Ostrowsky B, Greenko J, Adams E, Quinn M, O’Brien B, Chaturvedi V, Berkow E, Vallabhaneni S, Forsberg K, Chaturvedi S, Lutterloh E, Blog D, Group CaIW. 2020. Candida auris Isolates Resistant to Three Classes of Antifungal Medications - New York, 2019. MMWR Morb Mortal Wkly Rep 69:6–9.

21. Mukkada S, Kirby J, Apiwattanakul N, Hayden RT, Caniza MA. 2016. Use of Fungal Diagnostics and Therapy in Pediatric Cancer Patients in Resource-Limited Settings. Curr Clin Microbiol Rep 3:120–131.

22. Lockhart SR, Etienne KA, Vallabhaneni S, Farooqi J, Chowdhary A, Govender NP, Colombo AL, Calvo B, Cuomo CA, Desjardins CA, Berkow EL, Castanheira M, Magobo RE, Jabeen K, Asghar RJ, Meis JF, Jackson B, Chiller T, Litvintseva AP. 2017. Simultaneous Emergence of Multidrug-Resistant Candida auris on 3 Continents Confirmed by Whole-Genome Sequencing and Epidemiological Analyses. Clin Infect Dis 64:134–140.

23. Osei Sekyere J. 2018. Candida auris: A systematic review and meta-analysis of current updates on an emerging multidrug-resistant pathogen. Microbiologyopen 7:e00578.

24. Frias-De-Leon MG, Hernandez-Castro R, Vite-Garin T, Arenas R, Bonifaz A, Castanon-Olivares L, Acosta-Altamirano G, Martinez-Herrera E. 2020. Antifungal Resistance in Candida auris: Molecular Determinants. Antibiotics (Basel) 9.

25. Grahl N, Demers EG, Crocker AW, Hogan DA. 2017. Use of RNA-Protein Complexes for Genome Editing in Non-albicans Candida Species. mSphere 2.

26. Jensen-Pergakes KL, Kennedy MA, Lees ND, Barbuch R, Koegel C, Bard M. 1998. Sequencing, disruption, and characterization of the Candida albicans sterol methyltransferase (ERG6) gene: drug susceptibility studies in erg6 mutants. Antimicrob Agents Chemother 42:1160–7.

